# Adaptive Encoding of Coordinating Information in the Crayfish Central Nervous System

**DOI:** 10.1101/410951

**Authors:** Anna C. Schneider, Felix Blumenthal, Carmen R. Smarandache-Wellmann

## Abstract

Locomotion is essential for an animal’s survival. This behavior can range from directional changes to adapting the motor force to the conditions of its surroundings. Even if speed and force of movement are changing, the relative coordination between the limbs or body segments has to stay stable in order to provide the necessary thrust. The coordinating information necessary for this task is not always conveyed by sensory pathways. Adaptation is well studied in sensory neurons, but only few studies have addressed if and how coordinating information changes in cases where a local circuit within the central nervous system is responsible for the coordination between body segments at different locomotor activity states.

One system that does not depend on sensory information to coordinate a chain of coupled oscillators is the swimmeret system of crayfish. Here, the coordination of four coupled CPGs is controlled by central Coordinating Neurons. Cycle by cycle, the Coordinating Neurons encode information about the activity state of their home ganglion as burst of spikes, and send it as corollary discharge to the neighboring ganglia. Activity states, or excitation levels, are variable in both the living animal and isolated nervous system; yet the amount of coordinating spikes per burst is limited.

Here, we demonstrate that the system’s excitation level tunes the encoding properties of the Coordinating Neurons. Their ability to adapt to excitation level, and thus encode relative changes in their home ganglion’s activity states, is mediated by a balancing mechanism. Manipulation of cholinergic pathways directly affected the coordinating neurons’ electrophysiological properties. Yet, these changes were counteracted by the network’s influence. This balancing may be one feature to adapt the limited spike range to the system’s current activity state.

## 1. Introduction

One remarkable feature of animals is their ability to extract useful information from ubiquitous noise to ensure the organism’s functionality. For example, sensory neurons adapt to occurring ranges of stimulus intensities to maximize information transfer via their electrical activity (Dean et al., 2005; Laughlin, 1981; Maravall et al., 2007), a process in accordance with Barlow’s efficient coding hypothesis (Barlow, 1961). Most studies investigating neural adaptation and gain control focus on perception of environmental stimuli and adaptation of sensory neurons. Less is known about adaptive abilities of interneurons which provide information about internal states. Such interneurons can play important roles in coupling of neural networks that underlie behavior. Coupling networks through interneurons allows for faster information exchange compared to coupling via sensory pathways (LeGal et al., 2017). For example, lamprey increase their respiratory frequency in response to increased activity of the mesencephalic locomotor region even before proprioceptive neurons signal a higher energy demand (Gariépy et al., 2012). As such behavioral states (e.g. high or low locomotor activity) persist for some time the question arises if and how coupling interneurons adapt their encoding of state-related information.

In this study, the crayfish swimmeret system serves as model to examine adaptive encoding of coordinating information at different behavioral states (in this case low or high activity). The four pairs of pleopods (swimmerets) move in a metachronal wave from posterior to anterior during swimming, walking, posture control, or ventilation (Chrachri and Neil, 1993; Huxley, 1880; Mulloney and Smarandache-Wellmann, 2012; Neil and Miyan, 1986). Each swimmeret is controlled by its own microcircuit located in the respective abdominal hemiganglion (Figure 1A). Reciprocally inhibiting groups of non-spiking interneurons form a central pattern generator (CPG), which drives the motor neuron pools in alternating power-strokes (PS) and return-strokes (RS) (Mulloney, 2003; Paul and Mulloney, 1985a, 1985b; Smarandache-Wellmann et al., 2013). Two types of coordinating interneurons are present in each microcircuit: The Ascending (ASC_E_) and Descending (DSC) Coordinating Neuron. Both encode information about their home module’s activity state and send it as corollary discharge to anterior or posterior target ganglia, respectively. The target ganglia’s Commissural Interneuron 1 (ComInt 1, in figures C1) receives the coordinating information and conveys it to their module’s CPG (Mulloney and Hall, 2003; Smarandache et al., 2009; Smarandache-Wellmann et al., 2014).

**Figure 1:**
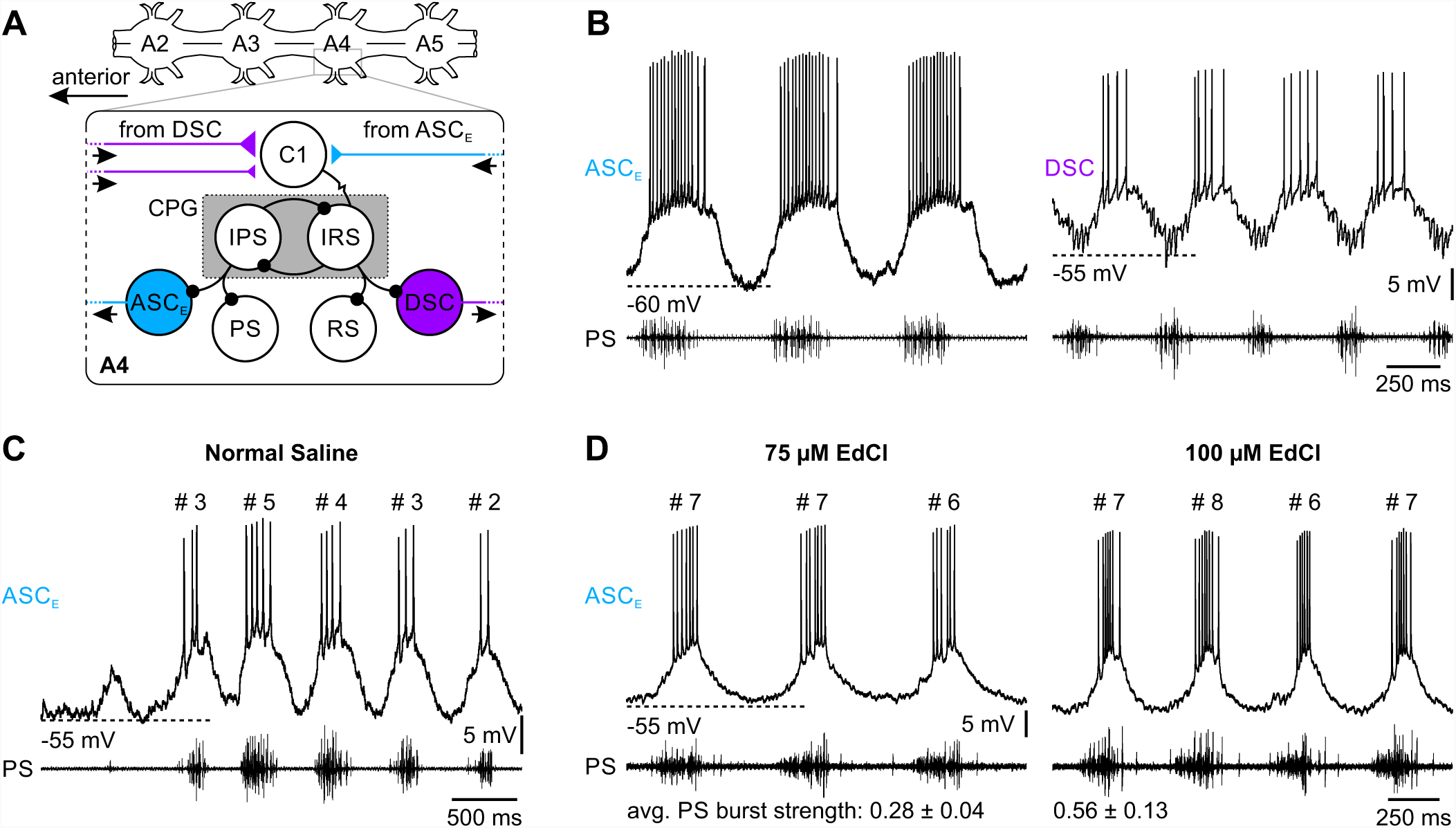
Encoding of coordinating information. **(A)** Schematic of the abdominal ganglia chain A2 to A5 with the microcircuit in the left hemiganglion of abdominal ganglion 4 (A4) that controls one swimmeret. The central pattern generator consists of mutually inhibiting IPS and IRS. IPS inhibits power-stroke (PS) motor neurons and ASC_E_ (cyan). IRS inhibits return-stroke (RS) motor neurons and DSC (purple). Coordinating information from the other three ipsilateral hemiganglia arrives via excitatory synapses at Commissural Interneuron 1 (C1). Inhibitory synapses are drawn as filled circles, excitatory synapses as triangles (size corresponds to synaptic strength). Arrows near coordinating neurons indicate direction of information propagation. **(B)** Activity of intracellularly recorded ASC_E_ and DSC (top trace) with their respective hemiganglion’s extracellularly recorded PS activity (lower trace). ASC_E_ is active in phase with PS, DSC in anti-phase to PS. **(C)** Variations in PS burst strength (approximation for size and frequency of active units) are tracked by ASC_E_’s spike count. **(D)** Preparations were locked with different edrophonium chloride (EdCl) concentrations at low or high excitation, indicated by average PS burst strength ± SD. At each excitation level changes in burst strength were tracked by the coordinating neuron’s spike count. However, across excitation levels the same number of spikes encoded different PS burst strengths.

The Coordinating Neurons couple the distributed CPGs and are necessary and sufficient to maintain the metachronal wave’s phase-lag of approximately 25% (Namba and Mulloney, 1999; Tschuluun et al., 2001). They receive the same input from the CPG as the swimmeret motor neurons (Smarandache-Wellmann and Grätsch, 2014). Hence, both neurons are oscillating time-locked to the motor neurons. ASC_E_ spikes in phase with PS, DSC spikes in anti-phase with PS (Figure 1B). PS burst strength, i.e. size and frequency of active motor units, is encoded by the number of spikes in ASC_E_ and DSC (Mulloney, 2005; Mulloney et al., 2006). In spontaneously occurring changes of PS burst strengths in the isolated preparation, a low spike count corresponds to weak, high spike count to strong motor activity (Figure 1C). Similar to *in vivo*, PS burst strength of the isolated abdominal ganglia chain can vary over a wide range in normal saline.

Application of excitatory neuromodulators, e.g. cholinergic agonists, can lock the system dose-dependent in low or high activity states (excitation levels) (Braun and Mulloney, 1993, 1995; Mulloney, 1997; Mulloney and Hall, 2007b). Surprisingly, at these forced excitation levels, the same number of coordinating spikes encodes different PS burst strengths in the same preparation, e.g. seven ASC_E_ spikes encode weak PS bursts at a low excitation level, but strong PS bursts at a high excitation level (Figure 1D). Additionally, forcing one part of the ganglia chain to a different excitation level than the other part results in changes in intersegmental coordination at the excitation boundary (Braun and Mulloney, 1995). This mismatch suggests that the system’s excitation level may tune the encoding properties of the coordinating neurons and/or the decoding abilities of ComInt 1. In this study, we present evidence that encoding of coordinating information adapted to the system’s excitation level both directly by altering intrinsic properties of coordinating neurons and indirectly via the network’s influence on these neurons. This provided a balancing mechanism for ASC_E_ and DSC to normalize their response range to occurring PS burst strengths, maintaining the rigorous coordination between CPGs. Furthermore, ComInt 1 reacted to similar changes in the excitation level, indicating that this postsynaptic neuron is tuned as well.

## 2. Materials and methods

All experiments were conducted on isolated abdominal ganglia chains of adult crayfish *Pacifastacus leniusculus*, DANA 1852, of both sexes. Animals were caught in the river Wupper at Müngstener Brückenpark, Germany, by a local fisherman and kept in freshwater tanks with hideouts at 14°C until sacrificing.

### 2.1 Dissection

The ventral nerve cord from thoracic ganglion (T) T4 to abdominal ganglion (A) A6 was dissected as described by Seichter and colleagues (2014). In brief, animals were anesthetized by chilling in ice for 20 min, exsanguinated with normal areated saline (in mM: 195.0 NaCl, 5.4 KCl, 2.6 MgCl_2_, 15.8 CaCl_2_, buffered with 1 M Trizma base and 0.5 M maleic acid at pH 7.4), and decapitated. The abdominal nerve cord was carefully dissected from the ventral pleon plate and pinned dorsal side up in a Sylgard-lined (Dow Corning, Midland, Michigan, USA) Petri dish. All ganglia were desheathed with fine scissors.

### 2.2 Electrophysiology

Fictive motor activity was recorded extracellularly with stainless steel differential electrodes from the posterior branch of the first segmental nerve N1, which carries axons of power-stroke (PS) motor neurons, of ganglia A2 – A5. The recording electrode and nerve were insulated from the bath; the reference electrode was placed in the bath nearby. All extracellular signals were amplified with a differential amplifier (MA102, Animal Physiology Electronics Lab, University of Cologne, Cologne, Germany). Signals of all recordings were digitized with a Digidata 1400 (Molecular Devices, Sunnyvale, CA, USA) or Micro1401-3 with Expansion ADC12 (CED, Cambridge, UK) at 10 kHz, and recorded with Clampex (Molecular Devices) or Spike2 (CED), respectively.

In order to identify Coordinating Neurons electrophysiologically, their activity was recorded with custom-made extracellular suction electrodes, which were placed in the appropriate locations on the Minuscule Tract (MnT) dorsal to the Lateral Giant axon (Mulloney et al., 2003; Namba and Mulloney, 1999). Signals from suction electrodes were preamplified 50-fold (MA103, Electronics Lab). To record intracellularly from the coordinating neurons and ComInt 1, sharp electrodes were pulled from borosilicate capillaries (o.d. 1.0 mm, i.d. 0.5 mm) on a P-1000 micropipette puller (Sutter Instruments, Novato, CA, USA) and filled with 1% dextran Texas Red (dTR; Invitrogen, Carlsbad, CA, USA) in 1 M KAc + 0.1 M KCl (electrode resistance 30 MΩ - 45 MΩ). Intracellular electrodes were connected to an intracellular amplifier SEC-05X (npi Electronic Instruments, Tamm, Germany). We recorded intracellularly from primary neurites of Coordinating Neurons in the Lateral Neuropil (LN) and identified them by matching the spikes on the intracellular recording to that of the extracellular suction electrode recordings, their phase-locked activity with the PS recordings, and their influence on the motor output of target ganglia upon de- or hyperpolarization (Jones et al., 2003; Mulloney and Hall, 2007a; Namba and Mulloney, 1999). The non-spiking ComInt 1 was recorded at its primary neurite at the ganglion’s midline. It can be identified by bursts of excitatory postsynaptic potentials (EPSPs) that are elicited by the other three ipsilateral Coordinating Neurons (Smarandache et al., 2009). Afterwards, the identity of the recorded neuron was confirmed by its morphology.

To set the system to different excitation levels, different concentrations of carbachol (in mM: 2, 3, 4; Sigma, St. Louis, MA, USA), or edrophonium chloride (EdCl; in mM: 50, 75, 100; Santa Cruz Biotechnology, Dallas, TX, USA) in 50 nM crustacean cardioactive peptide (CCAP; Bachem, Bubendorf, Switzerland) were perfused over the preparations. Carbachol- or EdCl-free saline was used for washing when switching excitation levels. Neurons were chemically isolated with low Ca^2+^ / high Mg^2+^ saline (low Ca^2+^ saline; same concentrations as normal saline except 0.6 mM Ca^2+^ and 16.4 mM Mg^2+^), and the excitation levels set with the same carbachol concentrations as mentioned above (Tschuluun et al., 2001).

To measure input resistance (R_in_), the neurons were hyperpolarized subthreshold to prevent spiking, and brief hyperpolarizing pulses (50 ms – 75 ms, -1 nA) were applied every 5 s (Figure 2A). R_in_ was measured at the same trough or membrane potential (V_m_) for each experiment.

**Figure 2:**
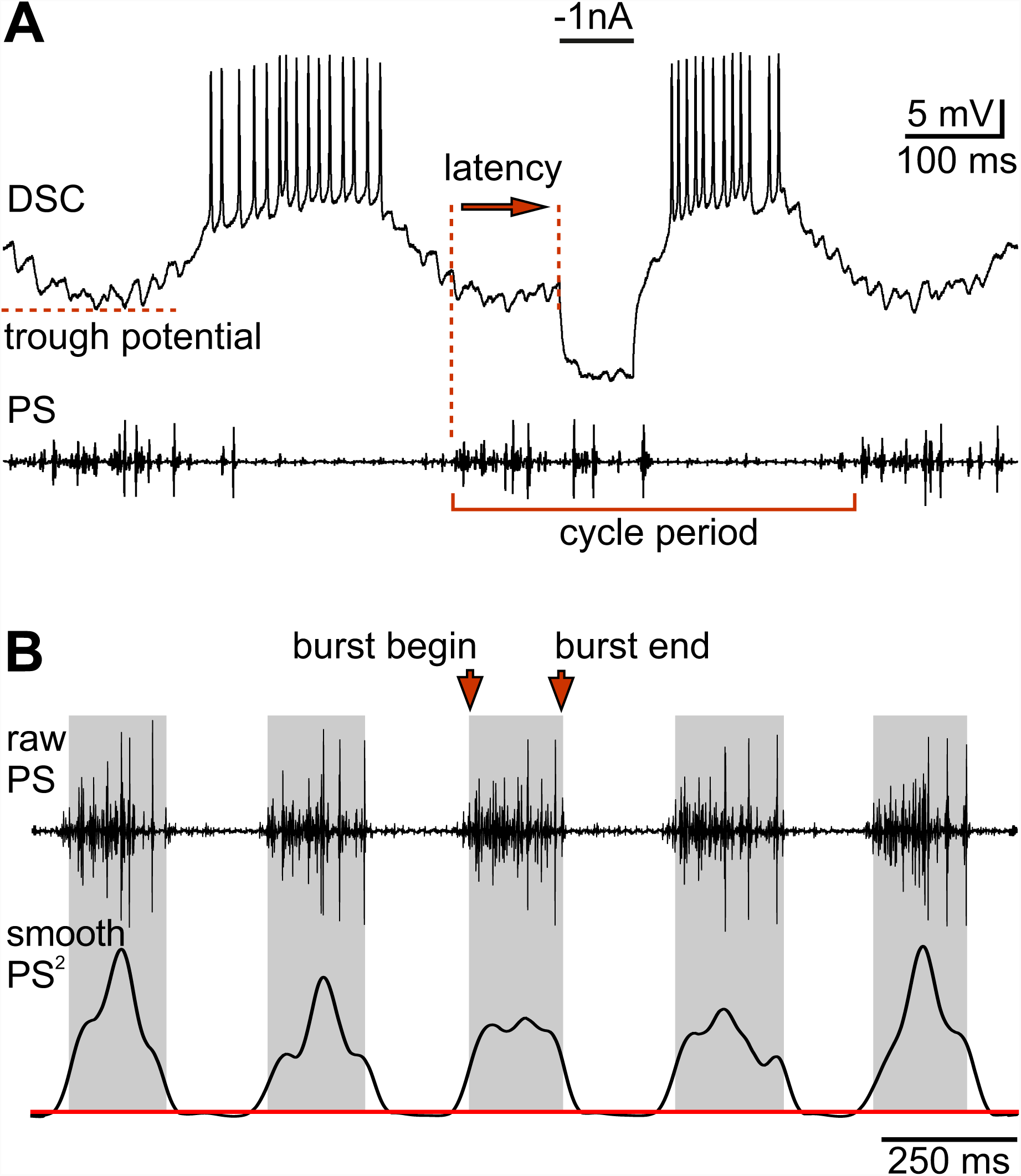
Analyzed parameters. **(A)** For each experiment and excitation level cycle period was measured as time from the beginning of a reference power-stroke (PS) burst to the beginning of the next. Input resistance was calculated by dividing membrane deflection by injected current. Phase of input resistance measurement was calculated by dividing the latency from PS burst begin to stimulus begin by cycle period. Trough potential was defined as most hyperpolarized membrane potential during each cycle. **(B)** To calculate PS burst strength the raw recording trace (top trace) was squared and smoothed (lower trace). Burst intensity is the area under the smoothed curve above noise threshold (red line) between burst begin and burst end (grey boxes, burst duration). Burst strength equals burst intensity divided by burst duration.

### 2.3 Data analysis

Cycle period and PS burst strength were determined for the hemiganglion in which a coordinating neuron was intracellularly recorded to characterize the system’s excitation level. Cycle period is the time from the beginning of a PS to the beginning of the next (Figure 2A). Burst strength is an approximation of the size and frequency of the overlapping active motor units during a PS burst (Figure 2B; (Mulloney, 2005)). To calculate burst strength, the PS trace was squared and smoothed with a Gaussian window kernel. The area under the smoothed curve between PS burst begin and end divided by burst duration is burst strength.

Burst strengths of individual experiments were normalized by the respective experiment’s maximum burst strength.

To investigate the electrophysiological properties of coordinating neurons at different excitation levels, we determined the neurons’ V_m_ (i.e. trough potential when they were oscillating), number of spikes per burst, and R_in_ by dividing the membrane deflection by the amplitude of the injected hyperpolarizing current (Figure 2A). Raw data of R_in_ measurements are lowess smoothed for easier visualization.

A full analysis of ComInt 1’s decoding properties is beyond the scope of this paper. Hence, we only analyzed changes in V_m_ and oscillation amplitude when we applied 3 µM carbachol.

N denominates the number of animals used; n denominates the trials per animal. Values are given as mean ± standard deviation, or median. For statistical analysis, either paired t-tests or Wilcoxon rank sum / signed rank tests were performed.

## 3. Results

In this study we focused on the adaptive encoding of information using the crayfish swimmeret system as example. The system’s excitation level can be set with different chemical concentrations. At all excitation levels, the swimmerets are still rigorously coordinated in a metachronal wave with approximate 25% phase-lag between segments. This phase-lag is independent of the rhythm frequency (Braun and Mulloney, 1993; Mulloney, 1997). In the following we will illustrate changes in the motor output at different excitation levels and in the Coordinating Neurons, which are necessary and sufficient for the proper coordination of the motor activity. Because bath application of chemicals can influence all neurons in a network, we investigated the Coordinating Neurons with both all synapses intact and chemically isolated by blocking transmitter release with low Ca^2+^ / high Mg^2+^ saline (Tschuluun et al., 2001). When the neurons were isolated, Coordinating Neurons and motor neurons became tonically active or silent and the swimmeret rhythm was no longer present.

### 3.1 System excitation

25 years ago the cholinergic agonist carbachol was established to activate and modulate the motor output of the isolated crayfish abdominal nerve cord, because it combines the effect of muscarinic and nicotinic agonists (Braun and Mulloney, 1993). Muscarinic agonists induce rhythmic activity in a quiescent preparation. Nicotinic agonists do not activate quiescent preparations but strongly modulate the motor output of already active preparations in a dose-dependent fashion. Similar modulation can be achieved by the application of acetylcholine esterase inhibitors like EdCl. Higher concentrations excite the system to higher levels (Mulloney, 1997; Namba and Mulloney, 1999). If preparations used for EdCl experiments did not show spontaneous rhythmic activity, we induced it with 50 nM CCAP.

To quantify which motor output characteristics change when changing the excitation level, we recorded PS activity at different carbachol or EdCl (+CCAP) concentrations. When the concentrations increased, cycle period decreased in four of five carbachol experiments on average by 94 ms ± 41 ms between 2 µM and 4 µM (N = 5, Figure 3Ai, paired t-test p < 0.021) and in all EdCl experiments by an average of 232 ± 112 ms between 50 µM and 100 µM (N = 6, Figure 3Aii, paired t-test p < 0.006).

**Figure 3:**
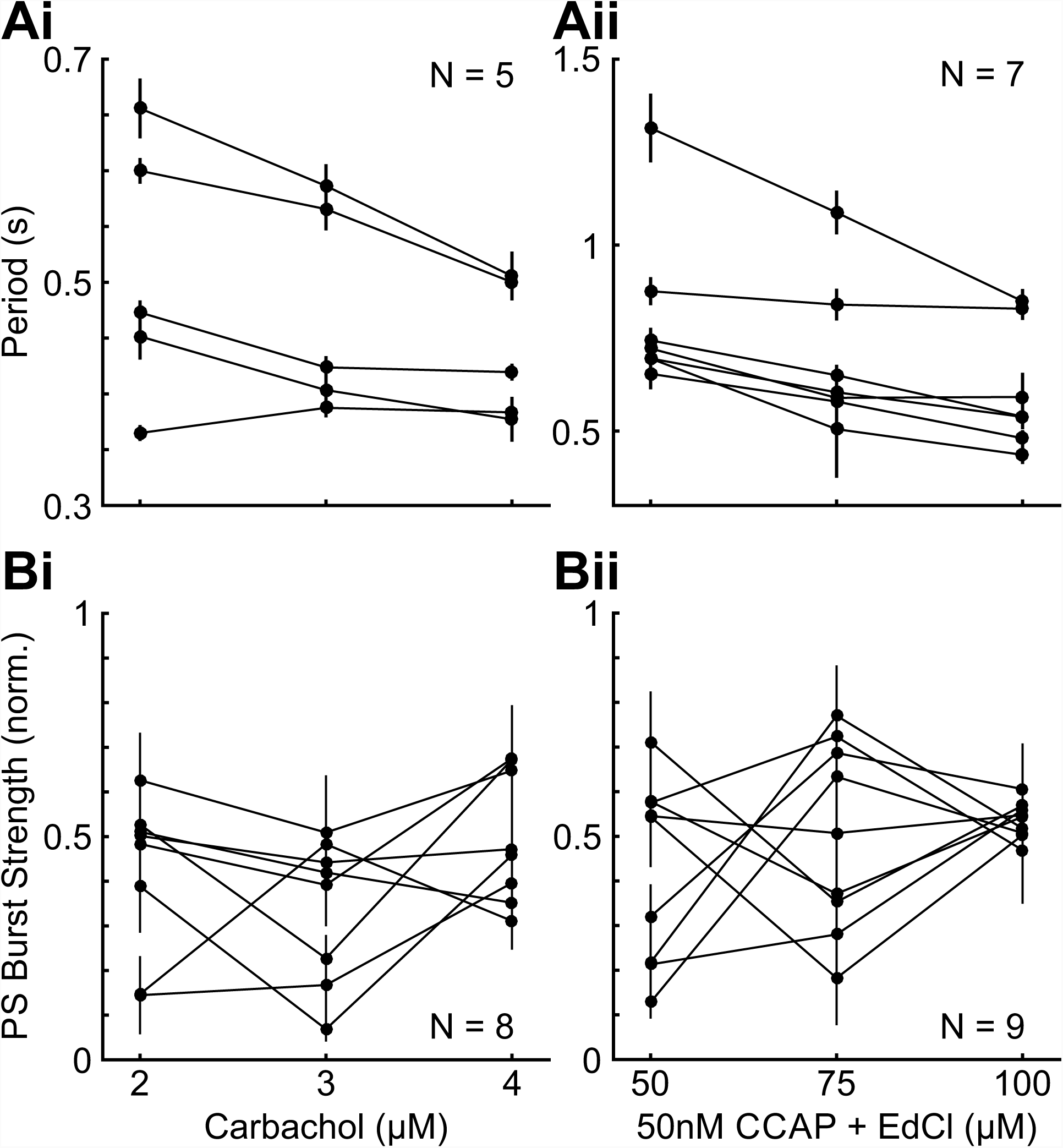
Effect of cholinergic modulation on cycle period and burst strength. **(A)** Period shortened with increasing carbachol concentrations in 4 of 5 experiments (Ai); two-tailed paired t-test p < 0.021. Similarly, period shortened in increasing edrophonium chloride (EdCl) concentrations in 7 of 7 experiments (Aii); two-tailed paired t-test p < 0.006. **(B)** Power-stroke (PS) burst strength was not as correlated to carbachol (Bi; N = 8) or EdCl (Bii; N = 9) concentrations as period. CCAP: Crustacean cardioactive peptide. Values are mean ± SD. Individual experiments are connected by lines.

Both chemicals did not have such a clear dose-dependent effect on PS burst strength (Figure 3B). Depending on the experiment, increasing the concentration had differential effects.

Varying the concentrations changed burst strength significantly in all experiments, although not strictly dose-dependent. However, in about two-thirds of the experiments (10 of 17) PS burst strength was higher in the highest concentration compared to lowest concentration.

As burst strength is an approximation of unit size and frequency in a burst, higher burst strength means that larger units are recruited and units are spiking at a higher frequency. Because a system’s excitation level corresponds to the excitability of its neurons, PS burst strength is a better representation of the excitation level than cycle period.

### 3.2 Adaptive encoding of coordinating information at different excitation levels

Previous experiments on the coordination of the swimmeret rhythms have shown that the Coordinating Neurons encode motor burst strength of their home ganglion by the number of spikes in their own bursts (Mulloney et al., 2006; Smarandache-Wellmann and Grätsch, 2014). Mulloney and colleagues (2007b) observed that this linear relationship does not hold when artificially changing the system’s excitation level. Hence, we calculated the average PS burst strength for any given number of spikes in the Coordinating Neurons at three different carbachol or EdCl concentrations (Figure 4). This revealed that, for each excitation level, the spontaneous variation in burst strength was still tracked by the coordinating neurons. Across excitation levels, however, the same number of spikes could encode for different burst strengths. Thus, the coordinating neurons encoded relative burst strength and the excitation level shifted the “tuning curves” by gain rescaling.

**Figure 4:**
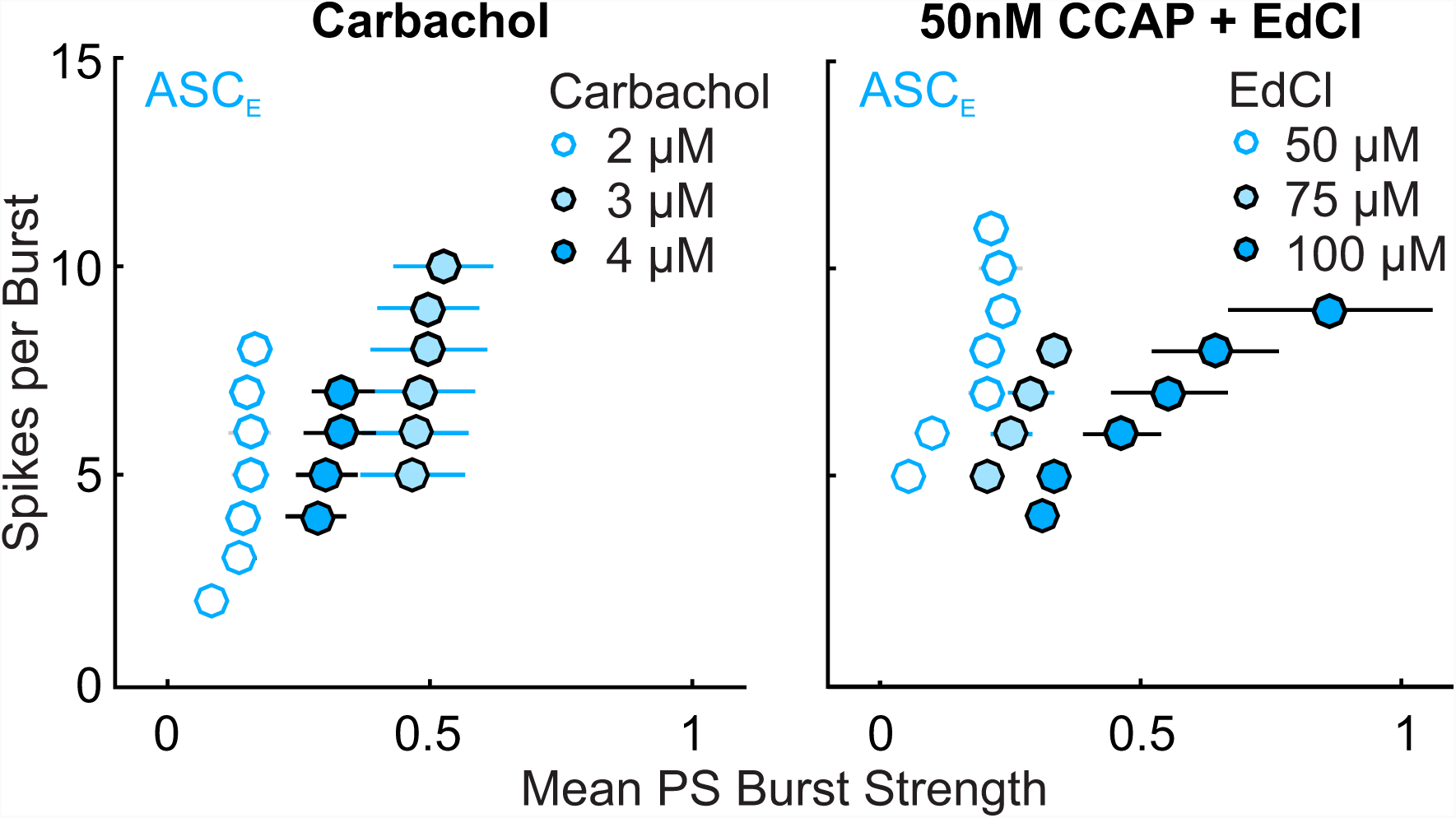
Adaptive encoding in ASC_E_. Exemplary data for one experiment in carbachol (n < 2000) and one experiment in edrophonium chloride (EdCl; n < 210). For each excitation level, indicated by different carbachol or EdCl concentrations, ASC_E_ tracked the changes in power-stroke (PS) burst strength. Across excitation levels the same number of ASC_E_ spikes could encode different PS burst strengths by shifting the tuning curve. The same pattern was demonstrated in 6 of 7 carbachol and 4 of 6 EdCl experiments. Values are mean ± SD.

### 3.3 Network effects stabilized the membrane potential

How are the Coordinating Neurons able to adapt their output to the system’s excitation level? To answer this question, we investigated their electrophysiological properties at different excitation levels. First, we examined the Coordinating Neurons’ V_m_ under the influence of different carbachol or EdCl concentrations (Figure 5). When all synapses were functional, DSC’s trough potential did not change (0 mV ± 4.5 mV) between lowest and highest chemical concentration (N = 8, Wilcoxon not significant; Figure 5C). ASC_E_’s V_m_ depolarized by 2.4 mV ± 2.5 mV (N = 10, Wilcoxon p = 0.02; Figure 5A, C). In contrast, when the neurons were synaptically isolated, ASC_E_ depolarized by 12.4 mV ± 7.1 mV (N = 11, Wilcoxon p < 0.001; Figure 5B, C), and DSC by 12.0 mV ± 4.8 mV (N = 8, Wilcoxon p < 0.008; Figure 5C). This illustrates that the chemicals we used to set the excitation level acted on the Coordinating Neurons both directly, causing the depolarization in the synaptically isolated condition, and indirectly via the network, causing the stabilization of V_m_ in the synaptically intact condition.

**Figure 5:**
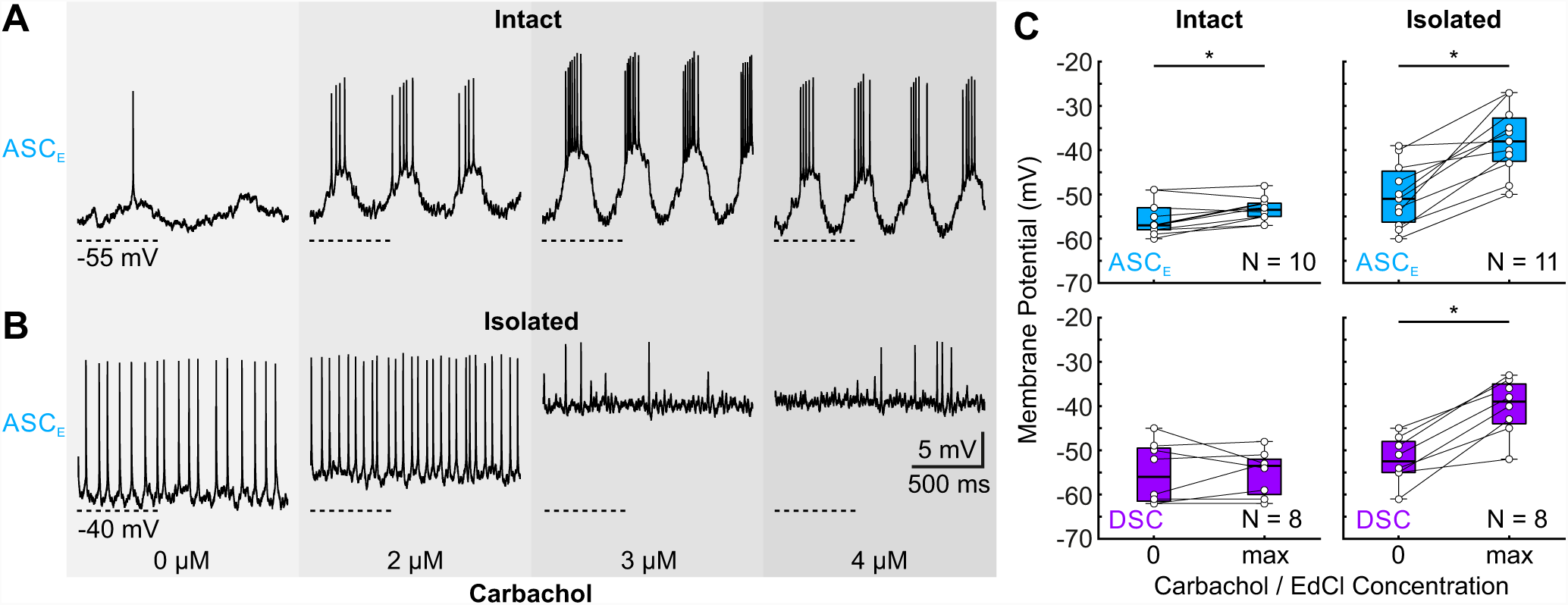
Differential effect of carbachol and EdCl on the membrane potential. **(A)** One example of the intracellular recording of ASC_E_ at different carbachol concentrations (grey shaded boxes) when all synapses were functional. The membrane potential did not change. **(B)** One example of ASC_E_ at different carbachol concentrations when synaptically isolated. The membrane potential depolarized with increasing carbachol concentrations. **C:** Membrane potentials for all ASC_E_s and DSCs at lowest and highest carbachol or edrophonium chloride (EdCl) concentration when synaptically connected (left plots; ASC_E_: N = 10, DSC: N = 8) or isolated (right plots; ASC_E_: N = 11, DSC: N = 8). In all isolation experiments the coordinating neurons’ membrane potential depolarized with higher chemical concentrations (Wilcoxcon signed rank test p < 0.001).

One possibility for the network to counteract the depolarization of the Coordinating Neurons would be to increase their inhibition by the CPG neurons. As these are non-spiking, already slight depolarization of their V_m_s would increase the overall release of inhibitory transmitter (Mulloney and Hall, 2007b). Therefore, we investigated the Coordinating Neuron’s input resistance (R_in_; Figure 6). In each cycle, ASC_E_’s and DSC’s R_in_ was highest when the neurons were producing spikes and lowest in their hyperpolarized phase (Figure 6A). Both neurons are inhibited by the non-spiking CPG neurons during their interbursts, which explains the low R_in_ during this phase (Smarandache-Wellmann and Grätsch, 2014). Applying different carbachol or EdCl concentrations shifted the R_in_ curves of both neurons (Figure 6A). As we were interested in how the neuronal properties changed at different excitation levels, and excitation level was not correlated to the chemical’s concentration in about one third of experiments, we compared a neuron’s R_in_ at that concentration which resulted in the average lowest PS burst strength (lowest excitation level) with the R_in_ at that concentration which resulted in the average highest PS burst strength (highest excitation level; Figure 6B). When the network was synaptically intact, ASC_E_’s R_in_ significantly decreased in 10 of 13 experiments (Wilcoxon p < 0.02 for individual experiments) and increased in the remaining three (Wilcoxon p < 0.001 for individual experiments). Similarly, DSC’s R_in_ decreased in 4 of 7 experiments (Wilcoxon p < 0.001 for individual experiments), increased in 2 of 7 experiments (Wilcoxon p < 0.001 for individual experiments), and did not change in 1 of 7 experiments. Hence, the Coordinating Neurons’ R_in_ decreased in the majority of experiments when the system’s excitation increased. As V_m_ was stable across excitation levels when the network was synaptically intact, this change in R_in_ must come from stronger inhibition to the Coordinating Neurons, presumably from the CPG. This increased inhibition could limit the amount of spikes these neurons can generate during a burst, keeping the spike count in a fixed range and adapting it to the system’s excitation level.

**Figure 6:**
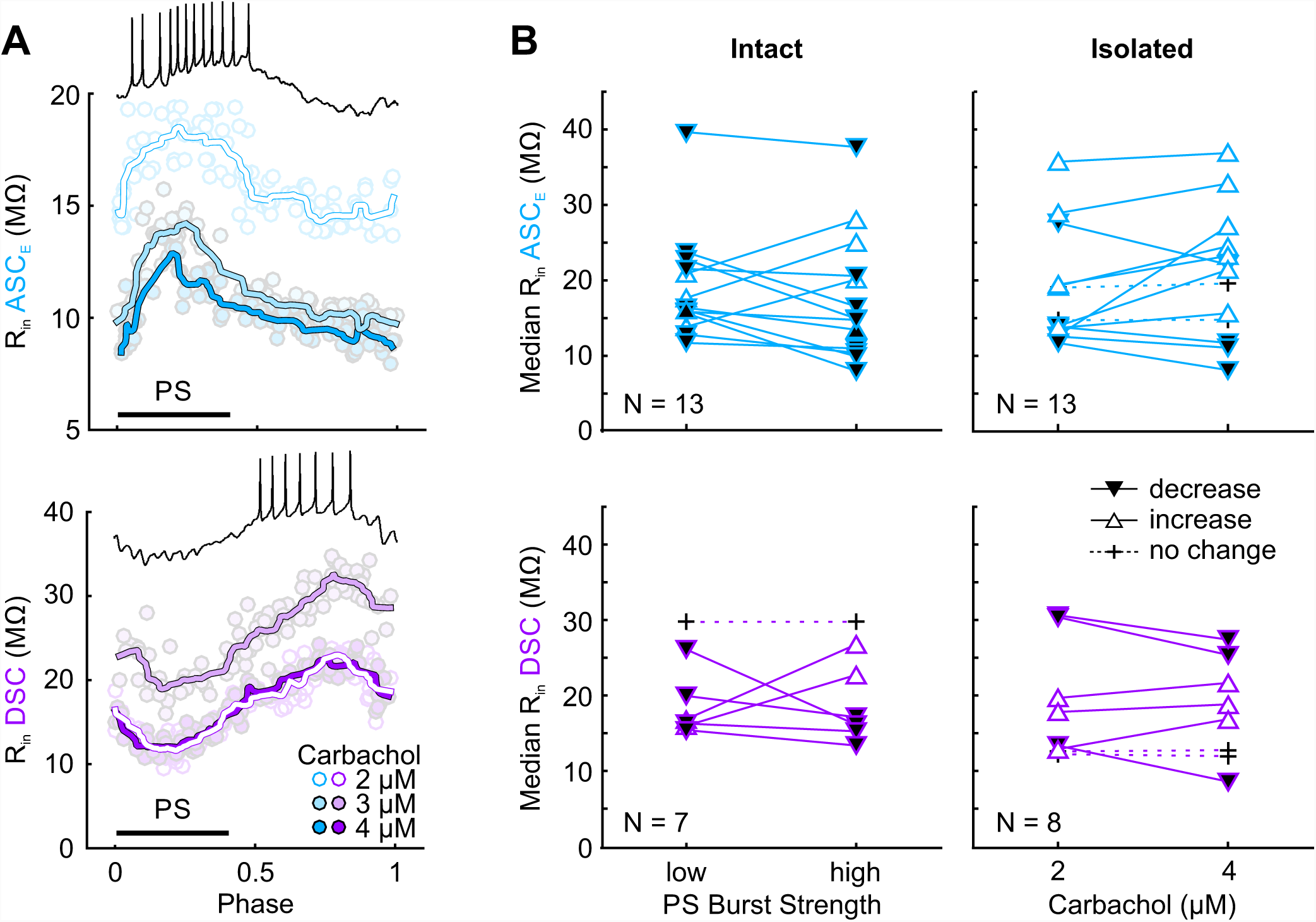
Differential effect of carbachol and EdCl on the input resistance. **(A)** One example of data for input resistance (R_in_) of ASC_E_ (top panel) and DSC (lower panel) during one cycle of activity at different carbachol concentrations. Power-stroke (PS) activity is indicated by black bar. Inset shows activity of the coordinating neurons during one cycle. Dots are individual R_in_ measurements. Line overlays are lowess smoothed measurements. **(B)** Median input resistance of all experiments compared between low and high excitation level indicated by mean PS burst strength in the synaptically connected condition (intact, left panels) or carbachol concentration in the synaptically isolated neurons (right panels). Black triangles indicate significant decrease in input resistance between low and high excitation level; Wilcoxon signed rank p < 0.05 for individual experiments. White triangles indicate significant increase in input resistance; Wilcoxon signed rank p < 0.05 for individual experiments. Crosses with dashed lines indicate no statistical difference in input resistance.Individual experiments are connected by lines.

Interestingly, the isolated ASC_E_’s R_in_ changed in the opposite direction when high carbachol concentrations were applied and ASC_E_ was clamped to its control V_m_. Please note that PS burst strength could not be calculated for isolated conditions. ASC_E_’s R_in_ increased in 7 of 13 experiments (Wilcoxon p < 0.001 for individual experiments), decreased in 4 of 13 experiments (Wilcoxon p < 0.001 for individual experiments), and remained unaltered in the remaining 2 experiments. The effect on DSC was not as pronounced. Its R_in_ increased in 3 of 8 experiments (Wilcoxon p < 0.001 for individual experiments), decreased in another three experiments (Wilcoxon p < 0.001 for individual experiments), and did not change in 2 experiments. Hence, the increased R_in_ correlated with the depolarization of the isolated ASC_E_ and DSC. This differential effect between isolated and intact conditions further emphasizes the balancing mechanism that acts by influencing the coordinating neurons directly and indirectly via the network.

### 3.4 Changes in excitation affect ComInt 1

Since the coordinating neurons adapt to the excitation level it is logical to assume that the postsynaptic receiver of coordinating information, ComInt 1, could be influenced by the system’s excitation level as well. To gain some first insights, we investigated its changes inV_m_ when changing the excitation level (Figure 7).

**Figure 7:**
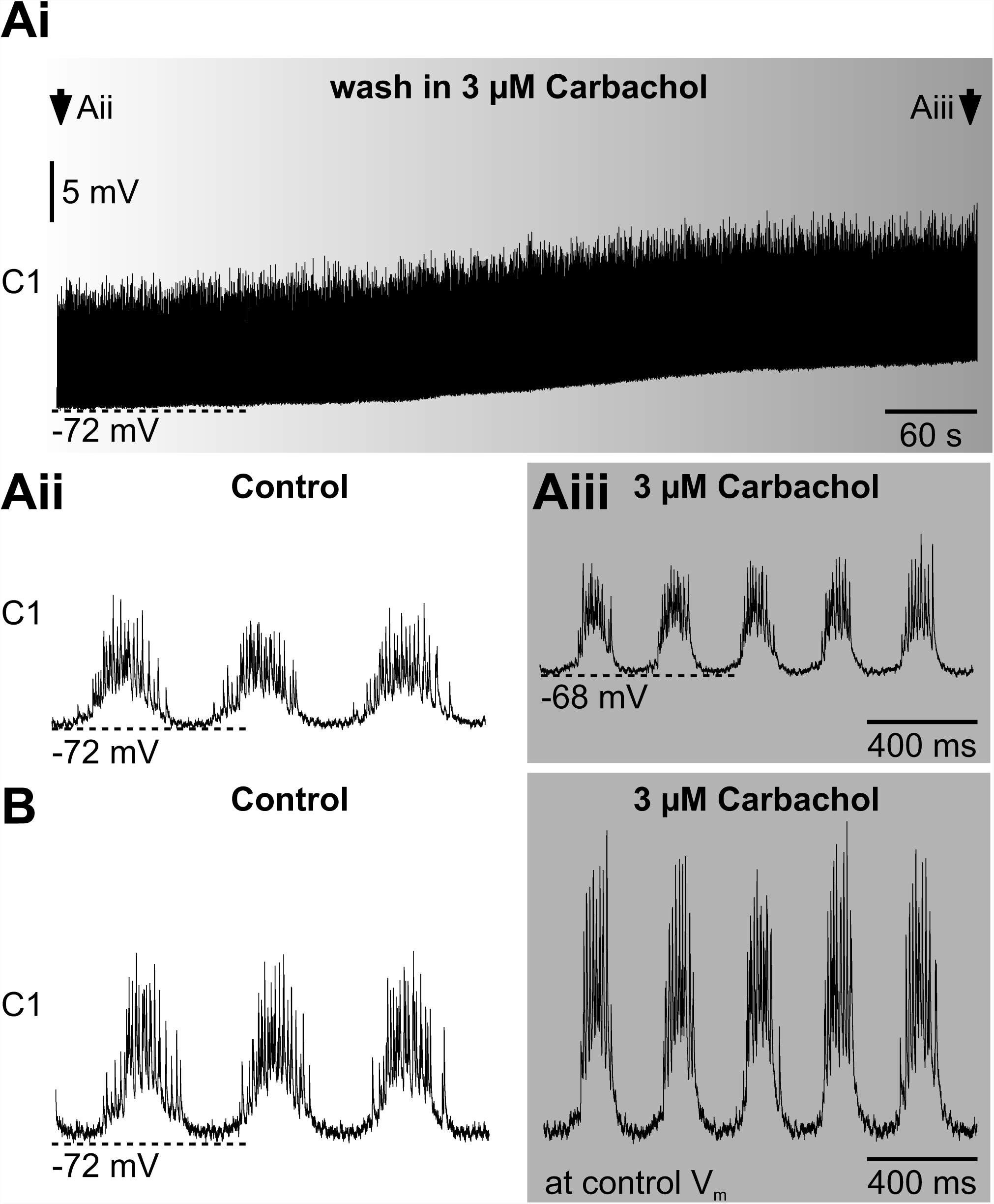
Carbachol depolarized ComInt 1. All recordings are from the same ComInt 1. **(Ai)** ComInt 1 (C1) depolarized during the wash in of 3 µM carbachol, indicated by the graded background fill. **(Aii)** ComInt 1 recording in normal saline at the time indicated by the left arrow in Ai. **(Aii)** ComInt 1 recording in 3 µM carbachol at the time indicated by the right arrow in Ai. **(B)** When ComInt 1 was hyperpolarized to the control trough potential in normal saline, both oscillation amplitude and EPSP amplitude increased.

In contrast to ASC_E_ and DSC, ComInt 1 depolarized with carbachol application (7 of 9 experiments; Figure 7 Ai, Aii). Yet, during this depolarization, the oscillation amplitude did not change in the majority of experiments (5 of 7 experiments; Figure 7 Aii), and decreased in 2 experiments. ComInt 1’s oscillation amplitude and EPSP amplitude increased when it was hyperpolarized to the same V_m_ as before carbachol application, (7 of 7 experiments; Figure 7 Aiii). These results show that the decrease in driving force for positive ions by ComInt 1’s depolarization was counteracted by a general increase in ComInt 1’s excitability. V_m_, oscillation amplitude and EPSP amplitude returned to control values when carbachol was washed out.

We show here that the same substances that affect the system’s excitation level and influence membrane properties of the coordinating neurons also induced changes in ComInt 1’s V_m_, indicating that this postsynaptic neuron is similarly tuned by the excitation level so that it can correctly interpret the arriving coordinating information.

## 4. Discussion

We have demonstrated that the Coordinating Neurons of the crayfish swimmeret system adapted to the system’s excitation level. The underlying mechanism for gain rescaling was influenced both by intrinsic properties of the Coordinating Neurons and by the network’s activity. Thus, the Coordinating Neurons were able to encode relative PS burst strength, which is correlated to the system’s excitation level. This effect is less pronounced in DSC, compared to ASC_E_. Because of the gradient of synaptic strength for arriving coordinating information in ComInt 1, ASC_E_’s impact is stronger than DSC’s (Smarandache et al., 2009). However, modeling studies have shown that the DSC input from the immediate neighboring ganglion is crucial to maintain the phase lag. With only long-range DSC input, phase lags decreased (Spardy and Lewis, 2018).

According to Barlow’s efficient coding hypothesis (Barlow, 1961), neurons adapt their output to represent a stimulus’ local distribution. This has been shown experimentally for the first time in the blowfly visual system (Laughlin, 1981). Here, the neurons’ output adapted to the stimulus’ statistical distribution that changed over time. Adaptive neurons conserve the amount of information transmitted per spike, and their coding accuracy is highest in the range of high stimulus probability (Dean et al., 2005; Maravall et al., 2007). In our study, the stimulus probability, i.e. PS burst strength, was determined by the system’s excitation level. Although it has never been thoroughly investigated, the amount of spikes the coordinating neurons generate during each cycle appears limited to a certain range: on average 10-20 for ASC_E_ and 3-10 for DSC during regular rhythmic activity (Mulloney et al., 2006; Mulloney and Hall, 2007b; Namba and Mulloney, 1999). Thus, the Coordinating Neurons must optimally utilize their finite capacity for information transmission by adapting their output range to the occurring PS burst strength distribution. Hence, ASC_E_ and DSC are able to encode PS burst strength at any given excitation level by spike count and rescale their encoding properties accordingly if excitation levels, and therefore PS burst strength distribution, change.

In this study we provide evidence that a balancing mechanism exists, which enables the gain rescaling of Coordinating Neurons. When synaptically isolated, their V_m_ depolarized and R_in_ increased in contrast to the intact network, with stable V_m_ and decreased R_in_ when increasing the excitation level. This means that increased system excitation resulted in increased graded inhibition by the non-spiking CPG neurons to reduce excitability of ASC_E_ and DSC. A decreased R_in_ can lead to shunting of synaptic input because the increased conductance lowers efficacy of electrotonic propagation (Laurent, 1990). Hence, any additional input the Coordinating Neurons might receive is less effective at higher excitation levels compared to low excitation levels. In contrast, high R_in_ can amplify synaptic input (Economo et al., 2014) which could boost input to the Coordinating Neurons at low excitation levels. Following this train of thought, the input from the non-spiking CPG neurons modulates the excitability of the Coordinating Neurons. In addition, another possible mechanism is that intrinsic history effects affect the Coordinating Neurons’ excitability and thus support adaptation to PS burst strength ranges. This has to be investigated in further studies.

The depolarization of the isolated neurons could not arise from synaptic input. Hence, ASC_E_ and DSC must possess cholinergic receptors. As their only identified afferents are inhibitory from the CPG (Smarandache-Wellmann and Grätsch, 2014), this result admits the possibility of additional unidentified excitatory input to the Coordinating Neurons. Interestingly, the depolarization was correlated to an increase in R_in_. Hyperpolarization-activated cyclic nucleotide-gated cation (HCN) channels and voltage-dependent sodium channels close at depolarization. While this might add to the changes in R_in_ during each cycle of oscillating activity, involvement of these channels to increase R_in_ under our experimental conditions seems unlikely. R_in_ was measured at subthreshold V_m_ between -60 mV and -50 mV, depending on the preparation. In this range the HCN channels are still open.

Similar changes for and R_in_ have also been observed for crustacean walking leg motor neurons (Cattaert et al., 1994). There, the inactivation of a voltage-gated outward K^+^ current underlies an exclusively muscarine-induced, long-lasting depolarization. Similar conclusions were drawn from Corronc and Hue (1993) for the V_m_ depolarization in the cockroach giant interneuron upon application of muscarinic agonists. In vertebrates, a non-inactivating voltage-gated low threshold K^+^ current that is inhibited by stimulation of muscarinic ACh receptors (I_M_) has been described (Adams et al., 1982; Alaburda et al., 2002; Brown and Adams, 1980; Halliwell and Adams, 1982). As the necessary KCNQ2/3 channel subunits appeared during the divergence of extant jawless and jawed vertebrates, I_M_ proper cannot be present in invertebrates (Hill et al., 2008; Wang et al., 1998; Yus-nájera et al., 2003). However, in *C. elegans* KCNQ-like K^+^ channels, with kinetics similar to that of vertebrates, have been identified (Wei et al., 2005). I_M_’s properties hyperpolarize V_m_ in response to depolarization, or depolarize V_m_ in response to muscarinic inhibition. Hence, this current can play a role in regulating a neuron’s excitability. In the swimmeret system, cholinergic agonists activate and modulate the motor output (Braun and Mulloney, 1993; Mulloney, 1997) so that it is likely for an M-like current to play a further role in balancing the coordinating neurons excitability, and therefore adaptation, based on the system’s excitation level.

This tuning allows the encoding of coordinating information over a wide range of system excitation with a limited spike range. While we could already demonstrate that ComInt 1’s is influenced when changing the system’s excitation level, further studies are needed to quantify the tuning of ComInt 1 membrane properties to interpret the arriving coordinating information.

## 5. Data availability

Datasets are available on request. The raw data supporting the conclusions of this manuscript will be made available by the authors, without undue reservation, to any qualified researcher.

## 6. Abbreviations

ASC_E_: Ascending Coordinating Neuron
CCAP: Crustacean cardioactive peptide
ComInt 1, C1: Commissural Interneuron 1
CPG: Central pattern generator
DSC: Descending Coordinating Neuron
dTR: Dextran Texas Red
EdCl: Edrophonium chloride
N1: First segmental nerve
PS: Power-stroke
R_in_: Input resistance
RS: Return-stroke
V_m_: Membrane potential, trough potential

## 7. Acknowledgements

This research was supported by an Emmy Noether DFG grant SM 206/3-1 and University of Cologne Advanced Researcher Group Grant ZUK 81/1.

We thank the group “Edelkrebsverein” and Ingo Selbach who provided us with the experimental animals.

Part of this study has been included in ACS doctoral thesis (Schneider, 2017).

## 9. Author contributions

ACS: Conceptualization, Experiments, Visualization, Writing—original draft FB: Experiments, Analysis

CSW: Conceptualization, Experiments, Analysis, Visualization, Resources, Supervision, Funding acquisition, Writing—original draft

## References

Adams, P. R., Brown, D. A., and Constanti, A. (1982). M-currents and other potassium currents in bullfrog sympathetic neurones. J. Physiol. 330, 537–572.

Alaburda, A., Perrier, J.-F., and Hounsgaard, J. (2002). An M-like outward current regulates the excitability of spinal motoneurones in the adult turtle. J. Physiol. 540, 875–881. doi:10.1113/jphysiol.2001.015982.

Barlow, H. B. (1961). “Possible Principles Underlying the Transformations of Sensory Messages,” in Sensory Communication, ed. W. A. Rosenblith (Cambridge: MIT Press), 217–234.

Braun, G., and Mulloney, B. (1993). Cholinergic modulation of the swimmeret motor system in crayfish. J. Neurophysiol. 70, 2391–2398.

Braun, G., and Mulloney, B. (1995). Coordination in the crayfish swimmeret system: differential excitation causes changes in intersegmental phase. J. Neurophysiol. 73, 880–885.

Brown, D. A., and Adams, P. R. (1980). Muscarinic suppression of a novel voltage-sensitive K+ current in a vertebrate neurone. Nature 283, 673–676. doi:10.1038/283673a0.

Cattaert, D., Araque, A., Buno, W., and Clarac, F. (1994). Nicotinic and muscarinic activation of motoneurons in the crayfish locomotor network. J. Neurophysiol. 72, 1622–1633.

Chrachri, A., and Neil, D. M. (1993). Interaction and synchronization between two abdominal motor systems in crayfish. J. Neurophysiol. 69, 1373–1383.

Corronc, H. L., and Hue, B. (1993). Pharmacological and Electrophysiological Characterization of a Postsynaptic Muscarinic Receptor in the Central Nervous System of the Cockroach. J. Exp. Biol. 181, 257–278.

Dean, I., Harper, N. S., and McAlpine, D. (2005). Neural population coding of sound level adapts to stimulus statistics. Nat. Neurosci. 8, 1684–1689. doi:10.1038/nn1541.

Economo, M. N., Martínez, J. J., and White, J. A. (2014). Membrane potential-dependent integration of synaptic inputs in entorhinal stellate neurons. Hippocampus 24, 1493–1505. doi:10.1002/hipo.22329.

Gariépy, J.-F., Missaghi, K., Chevallier, S., Chartré, S., Robert, M., Auclair, F., et al. (2012). Specific neural substrate linking respiration to locomotion. Proc. Natl. Acad. Sci. U. S. A. 109, E84–E92. doi:10.1073/pnas.1113002109.

Halliwell, J. V., and Adams, P. R. (1982). Voltage-clamp analysis of muscarinic excitation in hippocampal neurons. Brain Res. 250, 71–92. doi:10.1016/0006-8993(82)90954-4.

Hill, A. S., Nishino, A., Nakajo, K., Zhang, G., Fineman, J. R., Selzer, M. E., et al. (2008). Ion Channel Clustering at the Axon Initial Segment and Node of Ranvier Evolved Sequentially in Early Chordates. PLOS Genet. 4, e1000317. doi:10.1371/journal.pgen.1000317.

Huxley, T. H. (1880). The Crayfish?: An Introduction to the Study of Zoology. 1st ed. London: C. Kegan Paul & Co.

Jones, S. R., Mulloney, B., Kaper, T. J., and Kopell, N. (2003). Coordination of Cellular Pattern-Generating Circuits that Control Limb Movements: The Sources of Stable Differences in Intersegmental Phases. J. Neurosci. 23, 3457–3468.

Laughlin, S. (1981). A Simple Coding Procedure Enhances a Neuron’s Information Capacity. Z. Für Naturforschung C 36, 910–912. doi:10.1515/znc-1981-9-1040.

Laurent, G. (1990). Voltage-dependent nonlinearities in the membrane of locust nonspiking local interneurons, and their significance for synaptic integration. J. Neurosci. 10, 2268–2280.

LeGal, P., Dubuc, R., and Smarandache-Wellmann, C. R. (2017). “Interactions between rhythmic movement generators,” in Neurobiology of Motor Control: Fundamental Concepts and New Directions, eds. S. Hooper and A. Büschges (Wiley).

Maravall, M., Petersen, R. S., Fairhall, A. L., Arabzadeh, E., and Diamond, M. E. (2007). Shifts in Coding Properties and Maintenance of Information Transmission during Adaptation in Barrel Cortex. PLOS Biol. 5, e19. doi:10.1371/journal.pbio.0050019.

Mulloney, B. (1997). A Test of the Excitability-Gradient Hypothesis in the Swimmeret System of Crayfish. J. Neurosci. 17, 1860–1868.

Mulloney, B. (2003). During fictive locomotion, graded synaptic currents drive bursts of impulses in swimmeret motor neurons. J. Neurosci. 23, 5953–5962.

Mulloney, B. (2005). A method to measure the strength of multi-unit bursts of action potentials. J. Neurosci. Methods 146, 98–105. doi:10.1016/j.jneumeth.2005.01.020.

Mulloney, B., and Hall, W. M. (2003). Local commissural interneurons integrate information from intersegmental coordinating interneurons. J. Comp. Neurol. 466, 366–376. doi:10.1002/cne.10885.

Mulloney, B., and Hall, W. M. (2007a). Local and intersegmental interactions of coordinating neurons and local circuits in the swimmeret system. J. Neurophysiol. 98, 405–413. doi:10.1152/jn.00345.2007.

Mulloney, B., and Hall, W. M. (2007b). Not by spikes alone: Responses of coordinating neurons and the swimmeret system to local differences in excitation. J. Neurophysiol. 97, 436–450. doi:10.1152/jn.00580.2006.

Mulloney, B., Harness, P. I., and Hall, W. M. (2006). Bursts of Information: Coordinating Interneurons Encode Multiple Parameters of a Periodic Motor Pattern. J. Neurophysiol. 95, 850–861. doi:10.1152/jn.00939.2005.

Mulloney, B., and Smarandache-Wellmann, C. R. (2012). Neurobiology of the crustacean swimmeret system. Prog. Neurobiol. 96, 242–267. doi:10.1016/j.pneurobio.2012.01.002.

Mulloney, B., Tschuluun, N., and Hall, W. M. (2003). Architectonics of crayfish ganglia. Microsc. Res. Tech. 60, 253–265. doi:10.1002/jemt.10265.

Namba, H., and Mulloney, B. (1999). Coordination of Limb Movements: Three Types of Intersegmental Interneurons in the Swimmeret System and Their Responses to Changes in Excitation. J. Neurophysiol. 81, 2437–2450.

Neil, D. M., and Miyan, J. A. (1986). Phase-Dependent Modulation of Auxiliary Swimmeret Muscle Activity in the Equilibrium Reactions of the Norway Lobster, Nephrops Norvegicus L. J. Exp. Biol. 126, 157–179.

Paul, D. H., and Mulloney, B. (1985a). Local interneurons in the swimmeret system of the crayfish. J. Comp. Physiol. A 156, 489–502. doi:10.1007/BF00613973.

Paul, D. H., and Mulloney, B. (1985b). Nonspiking local interneuron in the motor pattern generator for the crayfish swimmeret. J. Neurophysiol. 54, 28–39.

Schneider, A. C. (2017). Encoding of Coordinating Information in a Network of Coupled Oscillators. Dissertation. Cologne, Germany: Universität zu Köln. Available at: http://kups.ub.uni-koeln.de/7592/ [Accessed May 23, 2018].

Seichter, H. A., Blumenthal, F., and Smarandache-Wellmann, C. R. (2014). The Swimmeret System of Crayfish: A Practical Guide for the Dissection of the Nerve Cord and Extracellular Recordings of the Motor Pattern. J. Vis. Exp. 93, e52109. doi:10.3791/52109.

Smarandache, C. R., Hall, W. M., and Mulloney, B. (2009). Coordination of Rhythmic Motor Activity by Gradients of Synaptic Strength in a Neural Circuit That Couples Modular Neural Oscillators. J. Neurosci. 29, 9351–9360. doi:10.1523/JNEUROSCI.1744-09.2009.

Smarandache-Wellmann, C., and Grätsch, S. (2014). Mechanisms of Coordination in Distributed Neural Circuits: Encoding Coordinating Information. J. Neurosci. 34, 5627–5639. doi:10.1523/JNEUROSCI.2670-13.2014.

Smarandache-Wellmann, C., Weller, C., and Mulloney, B. (2014). Mechanisms of Coordination in Distributed Neural Circuits: Decoding and Integration of Coordinating Information. J. Neurosci. 34, 793–803. doi:10.1523/JNEUROSCI.2642-13.2014.

Smarandache-Wellmann, C., Weller, C., Wright, T. M., and Mulloney, B. (2013). Five types of nonspiking interneurons in local pattern-generating circuits of the crayfish swimmeret system. J. Neurophysiol. 110, 344–357. doi:10.1152/jn.00079.2013.

Spardy, L. E., and Lewis, T. J. (2018). The role of long-range coupling in crayfish swimmeret phase-locking. Biol. Cybern. 112, 305–321. doi:10.1007/s00422-018-0752-3.

Tschuluun, N., Hall, W. M., and Mulloney, B. (2001). Limb Movements during Locomotion: Tests of a Model of an Intersegmental Coordinating Circuit. J. Neurosci. 21, 7859–7869.

Wang, H.-S., Pan, Z., Shi, W., Brown, B. S., Wymore, R. S., Cohen, I. S., et al. (1998). KCNQ2 and KCNQ3 Potassium Channel Subunits: Molecular Correlates of the M-Channel. Science 282, 1890–1893. doi:10.1126/science.282.5395.1890.

Wei, A. D., Butler, A., and Salkoff, L. (2005). KCNQ-like Potassium Channels in Caenorhabditis elegans CONSERVED PROPERTIES AND MODULATION. J. Biol. Chem. 280, 21337–21345. doi:10.1074/jbc.M502734200.

Yus-nájera, E., Muñoz, A., Salvador, N., Jensen, B. S., Rasmussen, H. B., Defelipe, J., et al. (2003). Localization of KCNQ5 in the normal and epileptic human temporal neocortex and hippocampal formation. Neuroscience 120, 353–364. doi:10.1016/S0306-4522(03)00321-X.

